# From ashes to adaptation: the impact of wildfires on the diet of *Podarcis lusitanicus* revealed by DNA metabarcoding

**DOI:** 10.1101/2025.01.31.635878

**Authors:** Catarina Simões, Diana S. Vasconcelos, Raquel Xavier, Xavier Santos, Catarina Rato, D. James Harris

## Abstract

Fire has long been recognized as an important ecological and evolutionary force in plant communities, but its influence on animals, particularly regarding predator-prey interactions, remains understudied. This study focuses on the impact of wildfires on the diet of *Podarcis lusitanicus*, a lizard species inhabiting a fire-prone region in the Iberian Peninsula. Faecal samples were collected from 12 distinct locations in Northern Portugal, at sites that burned in two distinct years (2016 and 2022), and in sites unburned since at least 2016. *Podarcis lusitanicus* is a generalist arthropod predator with dietary flexibility. Given the turnover of arthropod species after fire, it is expected to find variations in diet caused by different fire histories, especially between unburnt and recently burnt sites. Using DNA metabarcoding approach, employing high-throughput sequencing, the study revealed that while prey richness remained unaffected by wildfire regime, significant shifts occurred in diet composition between more recently burned and unburned areas, demonstrating a likely influence on prey availability after a more recent fire. Specifically, we found that differences in diet composition between these two fire regimes were due to the presence of *Tapinoma* ants and jumping spiders (*Salticus scenicus*) in unburned areas, while these were absent in areas burned in 2022. Interestingly, diets in unburned areas and areas burned in 2016 showed no significant differences, highlighting the lizards’ adaptability and the habitat’s resilience over time. *Talitroides topitotum* was found in dominance in both burnt areas, suggesting that this species may be fire tolerant. In addition, families such as Cicadellidae and Noctuidae were found to be more associated with more recently burned areas. The use of DNA metabarcoding in this study was essential to provide a more detailed and accurate view of predator-prey interactions in ecosystems susceptible to fire, providing a better understanding of changes in prey consumption in these environments.

## Introduction

Wildfires are a significant ecological disturbance, shaping the structure and dynamics of ecosystems worldwide [1]. They impact approximately one third of the Earth’s land surface [2], driving changes in species composition, habitat availability, and ecosystem functions [3, 4]. While many ecosystems, such as the Mediterranean vegetation, have evolved traits to persist under specific fire regimes [5, 6], rapid changes in fire frequency and intensity pose critical challenges to species’ survival. Human activities, particularly those that increase fire recurrence, exacerbate these challenges, altering habitat conditions and threatening biodiversity [5, 7]. Moreover, wildfires can have a direct effect, such as animal migration and/or mortality, and indirect effects on individual organisms by changing resource availability (e.g. food and shelter) and habitat structure with fires also often leading to the creation of a mosaic of fragmented habitats [8, 9]. These changes in environment determine the temporal and spatial dynamics of the fauna and flora, both in the short and long term [9].

After a fire, the responses of faunal populations are likely to vary among species depending on the life history characteristics and time elapsed since the fire [8]. Wildfires rarely result in total mortality among vertebrates [10], especially for species adapted to cliffs and rocks, whose habitats typically remain intact after the fire [11, 12], or for those with effective dispersal or burrowing abilities [13]. However, post-fire conditions can represent an increased risk due to reduced shelter and resources, and greater exposure to predators [14, 15].

While some taxa show resilience to fire disturbance, others are highly sensitive [16]. Arthropods, particularly in their immobile larval stages, are generally vulnerable, with poor dispersers being at greater risk [17]. Many insects, due to their flight capacity, are able to quickly recolonize affected areas, while surviving arthropods may benefit from the post-disturbance environment through diminished competition or a reduced risk of predation [18, 19]. Ants, for instance, demonstrate remarkable resilience by relocating to unburned areas, including underground refuges, which minimizes fire-related mortality [20]. In general, omnivorous fauna, including predators, tend to be more resilient and recolonize disturbed areas more quickly, as they are not reliant on specific trophic resources, allowing them to adapt to shifting food availability [21] and focus on the most abundant and easily captured prey species [22]. Grasshopper species may also temporarily increase in abundance in recently burned plots, as their generalist and invasive traits, combined with strong dispersal capabilities, allow them to thrive in post-fire conditions [20].

Fire has shaped predator-prey relationships for thousands of years [23], but with the increasing human-induced changes to Earth’s ecosystems, the interaction between fire and these dynamics now often leads to unpredictable and potentially harmful consequences [22]. In that sense, diet studies using DNA metabarcoding of prey can be useful to understand changes in trophic interactions mediated by fire occurrence, since they allow rapid and accurate identification of the taxa consumed by a predator. This is especially relevant in the current scenario of increased wildfires, to understand the impact of these alterations on the prey communities and how these affect the predator populations [24].

Portugal periodically faces serious damages due to wildfires [25, 26]. Climatic conditions, such as episodes of high rainfall followed by dry periods, coupled with strong winds, significantly increase the likelihood of natural wildfires in this country [27-29], especially in Northern Portugal where most of the country’s burned areas are found [30]. Lacertid wall lizards of the genus *Podarcis* (Wagler 1830), are a group of reptiles that evolved and diversified along the Mediterranean Basin [31, 32]. They represent a principal herpetofauna element of Mediterranean ecosystems, playing an important ecological role in food chains [33]. The Lusitanian wall lizard *Podarcis lusitanicus* is endemic to the Iberian Peninsula (Fig 1), being widespread in northern Portugal, northwest Spain, and northeast to Picos de Europa [34], and whose demographic history was highly influenced by the Pleistocene glaciations [35]. *Podarcis lusitanicus* is a saxicolous, insectivorous, diurnal and small lizard that tends to use walls, logs and areas with less vegetation for thermoregulation, foraging and shelter [31]. Across its Portuguese range, this lizard has an irregular distribution, with populations located on open natural rocky outcrops and artificial stone walls surrounding agricultural fields [36]. As a result of this requirement for more open areas [37], *P. lusitanicus* seems to benefit from recurrently burned landscapes, dominated by granitic rocks, where lizard populations reach high effective sizes [38, 39]. Fire removes the vegetation and reveals rocky outcrops, creating new habitats, which favours population expansion, dispersal between metapopulations and the recruitment of migrants, increasing the population genetic diversity [27]. However, the high rainfall rate in northern Portugal leads to a fast vegetation recovery that is prone to become burned again [40]. For lizards living in areas with recurrent fires, fire-vegetation-recovery cycles cause prey availability to change, depending on the time since the last fire.

**Fig 1.**
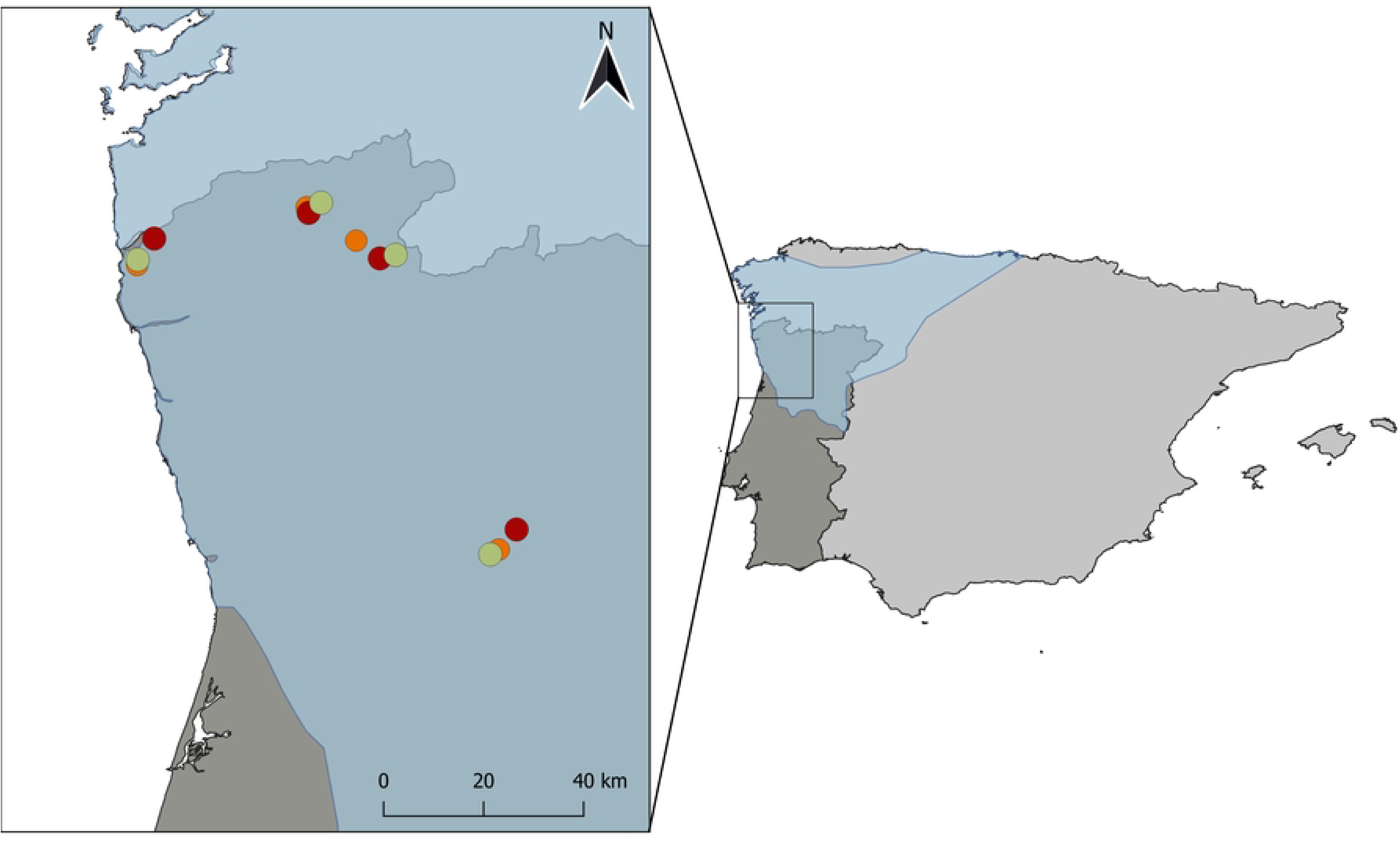
Location of the study sites in the NW extreme of Portugal. Each sampled location - Viana do Castelo (VC), Álvora (AL), Gerês-Soajo (GS) and Marco de Canaveses (MC) - is comprised by three sites with different historic fire conditions: unburned (UN in green), burned in 2016 (BU16 in orange), and burned in 2022 (BU22 in red). The approximate distribution range of *P. lusitanicus* is shown in blue, on the right map.

In this study, we assessed the impact of fires on the diet composition and diversity of the Lusitanian wall lizard based on a DNA metabarcoding approach. We described the diet of lizard populations living in recently burned plots (one year since the last fire), plots burned seven years since the last fire, and long-unburned plots. Given the high species turnover among arthropod groups after fire [16, 17, 41], we expect significant differences in the composition of lizards’ diets, especially between plots of the most different fire history.

## Material and Methods

### Study area

This study was performed in northwestern Portugal confined by the latitudes 41°58’ to 41°09’ N and longitudes -8°49’ to -7°56’ W (Fig. 1).

Four locations, Viana do Castelo (VC), Marco de Canaveses (MC), Gerês-Soajo (GS) and Álvora (AL), were chosen based on previous knowledge on habitat suitability and similarity (e.g. topography) and fire occurrence (ICNF; https://geocatalogo.icnf.pt/catalogo_tema5.html). At each location we selected an area burned in 2022 (coded as BU22), another burned in 2016 (coded as BU16), and a long-unburned area (coded as UN; no fires recorded since 2016 onwards), totalling 12 sampling sites (Fig 1), that include four replicates for each of the three fire occurrence groups (BU16, BU22 and UN). Selected areas representing BU16, BU22 and UN sites were in proximity with each other (< 10 Km radius) and showed similar overall landscapes, such as rocky habitats, as well as an identical vegetation structure and type between the sites with the same fire type, reducing the external variability of the landscape that could influence the presence or behaviour of the species.

### Sampling methods

Sampling of *Podarcis lusitanicus* lizards was carried out during the day (10 AM - 3 PM) between May and June 2023, across natural open rocky outcrops and artificial rock walls surrounding agricultural fields.

Lizards were captured with a noose. Generally, lizards defecated soon after capture, otherwise a simple abdomen massage was enough to obtain a freshly voided faecal sample. Faecal pellets were collected from 183 individuals of *P. lusitanicus*: 38 in Viana do Castelo, 50 in Marco de Canaveses, 53 in Álvora and 42 in Gerês-Soajo (S1 Table). All faecal samples were preserved in 96% ethanol.

Sex was determined by the presence of developed femoral pores, and the robustness of the head [42-44]. We also took standard morphological measurements, including weight and snout ventral length (SVL) using a digital scale and calliper, respectively. All measurements were taken by the same person (D.S. Vasconcelos) to eliminate inter-observer error. All individuals were released at the capture site on the same day.

### DNA Extraction and amplicon library preparation

The 183 pellet samples were extracted using the E.Z.N.A. Tissue DNA Kit (Omega Bio-Tek, U.S.A.), following the manufacturer’s instructions with a minor modification in the digestion step, using 800 μL of Gordon buffer instead of the 200 μL of TL buffer (from the kit) to improve DNA extraction of both hard and soft tissues [45]. All samples were vortexed to disrupt the faecal mass and digestion occurred over 3 hours. Extracted DNA was stored at -20°C. Eight extraction blank samples were also obtained to control for contaminants present in extraction kits and/or in the laboratory environment. A short fragment (∼205 bp) of the mitochondrial cytochrome c oxidase subunit I (COI) was amplified using Polymerase Chain Reaction (PCR) with the Fwh2 primers developed by Vamos et al. (2017) [46], which have been proposed as the most appropriate for the study of insectivorous diets [47, 48]. The primers were modified to include Illumina adaptors and a 0–5 bp addition of N bases between the adaptor and the primer to increase sequencing diversity and quality. The different primer variations were then combined before PCR reactions, resulting in mixed forward and reverse primer single solutions.

The PCR reaction consisted of 5 μL of QUIAGEN Multiplex PCR Master Mix (Quiagen, Crawley, UK), 0.3 μL combination of six Forward primers, 0.3 μL combination of Reverse Primers, 2.4 μL of ultra-pure water, and 2 μL of DNA extract. Cycling conditions used consisted of an initial denaturation at 95°C for 15 min, followed by 45 cycles of 95°C for 30s, annealing at 52°C for 45s, extension at 72°C for 20s, and a final extension at 60°C for 5min. Four PCR negative controls were used to detect possible contaminations in the PCR reagents.

An initial PCR clean-up to remove unused primers and primer dimers was performed, using the Agencourt AMPure XP beads (Beckman Coulter, Brea, CA, USA). This cleaning was performed using a proportion of 0.85 μL of magnetic beads to 1 μL of PCR product. To attach unique barcodes and Illumina sequencing indexes to each sample, a second PCR was performed using a distinct combination of index sequences per sample. This PCR index was performed using 2.8 μL of ultra-pure water, 7μL of 2x Kapa HiFi, 1.4 μL of each index, and 2.8 μL of cleaned DNA. Indexing the PCR required an initial denaturation of 95°C for 3min, followed by 10 cycles of 95°C for 30s, annealing at 55°C for 30s, extension at 72°C for 30s, and a final extension of 72°C for 5min. In order to confirm the success of the barcode’s incorporation in all samples, PCR products were tested in a 2% agarose gel with the *a priori* knowledge that amplicons should be ∼100 bp longer than in the previous PCR. A second PCR clean-up was then performed, under the same conditions as the first one, except that the beads’ ratio was 0.8.

After the second PCR clean-up, all indexed PCRs were quantified using Epoch™ Microplate Spectrophotometer (BioTek Instruments, Inc.; Winooski, VT, USA), followed by the normalisation to obtain the same concentration in all the samples. After the normalisation, the samples were pooled together. The pool was tested for quality control in a TapeStation 4200 High Sensitivity D1000 Assay (Agilent, Santa Clara, CA), and a cleaning was needed using a 0.78 ratio of beads. A second test for quality control in the TapeStation was performed to confirm the success of the cleaning. The final pool was sent to GENEWIZ Next Generation Sequencing laboratory with a final concentration of 27nM to be sequenced in an Illumina MiSeq sequencer with 2x250bp paired-end (PE) configuration and PhiX (≤30%) was used to increase sequencing diversity.

### Sequence denoising and taxonomic assignment

The software PEAR [49] was used to merge the forward and reverse reads into single sequences, discarding single and unassembled reads. Base pairs with less than 26bp of overlap were also rejected. Then, the primers were removed, using the command *ngsfilter* from OBITools [50]. The reads were once more counted to check for successful rates of the previous step with the *grep* command and then dereplicated into unique sequences using the *obiuniq* command. Lastly, sequence cleaning was performed to remove sequences with less than 10 reads and chimeras, using the *obiclean* command. Sequences with a length between 202bp and 208bp were kept and clustered at a 99% similarity threshold to form Operational Taxonomic Units (OTUs), using VSearch [51]. OTUs that were represented by a read count < 1% of the total number of reads obtained for each sample were removed [52]. This should allow the removal of most PCR and sequencing errors that still passed the *obiclean* denoising step. Additionally, all reads identified in the extraction and PCR controls were subtracted from the corresponding sample batch [53].

The obtained OTU table and sequences were further cleaned to remove redundant and unreliable OTUs using the R package LULU [54], and the taxonomic assignment of OTUs was performed using BOLDigger [55], followed by manual inspection and curation. We also used the BLAST function from NCBI implemented in Geneious Prime 2021.1.1 [56], when species-level assignments were not possible with BOLDigger. The taxonomic level assignment was performed according to the following thresholds: to the Class level if below 90%; to the Family level if between 90 and 95%; and to the Species or Genus level (in cases where more than one species from the same genus had a > 95% sequence match) if higher than 95% similarity.

Following the completion of taxonomic assignments, the distribution and presence of detected prey species in Portugal were verified using a combination of local databases, that included Naturdata (https://naturdata.com/) and Lusoborboletas (https://www.lusoborboletaspt.com/) and global databases, such as GBIF (http://www.gbif.org/). Additionally, relevant scientific articles were reviewed to confirm whether the presence of specific species within Portuguese ecosystems had already been documented.

### Statistical Analyses

Lizards body size (snout-vent length, hereafter SVL) and weight (hereafter BW) can influence diet composition [57, 58]. For this reason, we compared SVL and BW between sexes and among fire types. First, the normality of BW and SVL distributions was assessed using the Shapiro-Wilk test, which revealed that while SVL (p = 0.452) followed a normal distribution, BW did not (p = 0.0199). Consequently, both SVL and BW were log-transformed. Variation in logSVL and logWeight across sexes and fire types (burned in 2016, burned in 2022, and unburned) was visualized using boxplots. Body condition was analysed through a linear regression between logWeight and logSVL for males and females. An analysis of variance (ANOVA) was conducted using the *aov* function to examine sexual dimorphism in body size (logSVL) while accounting for Fire type and Sex as fixed factors, and locality as a random factor. Similarly, a separate ANOVA was conducted to analyse the variation in body mass (logWeight) using the same fixed and random factors. However, since logWeight and logSVL are correlated, subsequent analyses focused only on logSVL. Pairwise comparisons were carried out using the emmeans package 1.10.2 [59].

For the statistical analysis, R 4.1.0 [60] was used to assess differences in dietary descriptors (i.e., diet richness and composition). Dietary analysis was based on two different taxonomic levels: OTU (all taxonomic units identified to the most possibly resolved taxonomic level) and Family. For the OTUs not identified to the species level, we built a neighbour-joining tree in Geneious Prime v. 2024.0.2 (Biomatters), visually inspected the corresponding alignment, and checked for patterns of genetic similarity (∼98%) in order to cluster them into distinct taxa (e.g., Carabidae 1, Carabidae 2, and so on).

A Generalized Linear Mixed Model (GLMM) was implemented to investigate the effects of different predictors on diet richness (i.e. the number of different OTUs or Family prey types per lizard) while accounting for the replicate structure of the sampling scheme. This approach was chosen because the number of OTU or Family prey items per pellet followed a Poisson distribution, as determined by the Shapiro-Wilk test. Fire type, sex, and logSVL were included as fixed effects, while Locality was considered as a random factor. Pairwise comparisons were again carried out using the emmeans package. The magnitude (effect size) and ecological significance (positive or negative) of the predictor variables on OTU richness and Family richness were analysed by presenting the model coefficients and their respective 95% confidence intervals. When the confidence intervals include the value 0, this indicates that the effect is not statistically significant (p ≥ 0.05).

The iNEXT v. 3.0.1 package [61](Hsieh et al., 2016) was used to perform rarefaction and extrapolation curves to analyse whether the type of fire (unburned versus burned; burned in 2016 versus burned in 2022) and Sex had any impact on both the OTU and Family richness, setting the confidence level at 84% [62] with 1,000 bootstrap replicates. Further, using the *ggiNEXT* function both sample-based and coverage-based rarefaction curves were plotted. However, the estimated richness was compared considering completeness (i.e., sample coverage) instead of sample size (i.e., number of samples), to avoid biases of communities with different levels of richness requiring different sampling efforts to be sufficiently characterised [63].

Additionally, a rank abundance plot was created to visualise the top 10 most abundant prey species across the different Fire types, using the *ggplot* from the ggplot2 package in R [64].

To compare diet composition at the OTU and Family levels, a permutational multivariate analysis of variance (PERMANOVA) was used with Fire type, Sex and logSVL as fixed factors, using the R package vegan 2.6-6.1 [65], considering a stratified design (strata = Locality). The presence or absence of each prey item in each sample was used to build a Jaccard (among OTUs) and Bray-Curtis (among Families) similarity index using the *vegdist* function from the vegan package. After the PERMANOVA, a pairwise comparison was performed using the *pairwise.adonis2* function from the pairwiseAdonis R package [66].

A homogeneity of dispersion test (function *betadisper*) was performed to make sure that the differences observed with PERMANOVA were not due to unequally dispersed values across the different groups, as it evaluates the homogeneity of multivariate dispersions. Afterwards, a similarity percentage analysis was carried out, using the function *SIMPER* implemented in the vegan R package, to assess the contribution of each prey item to the differentiation between diets.

Finally, a redundancy analysis (RDA) was done using prey types (Families) with abundances ≥ 5 items using sex and Fire type as factors and locality as conditional to account for spatial heterogeneity. This analysis allows us to identify associations between specific prey Families with a particular fire type. The RDA was performed using the vegan package, and scaling was applied to standardize the data. A separate permutation test was run for each explanatory term to evaluate their individual contributions to the model.

## Results

The regression for logSVL and logweight shows that these parameters are significantly and positively correlated in both females and males (Females: r2 = 0.427; Males: r2 = 0.491), with males being larger and heavier than females (S1 Fig). The only parameter that affects SVL (p = 0.0116) is Sex, but the logWeight is significantly influenced by both Sex (p = 3.72e-05) and Fire type (p = 0.0201) (S2 Fig and S2 Table). The significant differences in weight obtained between the Fire types were due to differences between recently burned areas (BU22) and unburned ones (UN) (p = 0.0251).

### Sequence filtering

Approximately 23 million raw sequence reads were generated from the 183 faecal samples analysed. 357,866 reads were retained after quality control, resulting in 1,102 OTUs. Non-target amplification (37.39% of total reads) was observed from different sources, both in the samples and in the extraction and PCR negative controls. Most of the non-target OTUs were identified as belonging to Fungi (mainly Ascomycota and Basidiomycota), accounting for 12.53% of total reads and 33.51% of non-target OTU diversity. An expected presence of *Podarcis lusitanicus* (6.88% of total reads) was observed. The final dietary dataset comprised 237,402 reads, encompassing 537 distinct OTUs.

### Diet analyses

From a total of 537 prey items, five classes of Arthropoda (Arachnida, Collembola, Diplopoda, Insecta and Malacostraca), one class of Mollusca (Gastropoda), and one class of Annelida (Clitellata), were identified. These encompassed at least 26 distinct orders, 95 families and 93 species. In general, the order Araneae (21%) was the most consumed prey, with Salticidae being the most frequent Family (10%). The other most consumed prey belonged to Coleoptera (12%), Hemiptera (12%), Hymenoptera (11%) and Lepidoptera (7%) (S3 Fig).

The generalized linear mixed models (GLMM) showed that none of the variables had a significant effect on either OTU or Family diet richness (S3 Table). The estimated model coefficients and 95% confidence intervals (CIs) for OTU richness (S4A Fig) and Family richness (S4B Fig), indicate that the confidence interval for Fire type, Sex and LogSVL overlap 0, representing no significant effects among groups.

From the analysis of the rarefaction and extrapolation curves (Fig 2), we observed no difference between burned and unburned areas, at both the OTU and Family levels (Fig 2A). Likewise, areas burned in 2016 (BU16) and in 2022 (BU22) have overlapping niche breadths (Fig 2B). The same pattern is observed between sexes (Fig 2C). For all rarefaction analysis, the minimum observed sample coverage was always above 50%, except for the unburned areas in the OTU diversity analysis (Fig 2A) and the Females (F) in the OTU diversity analysis (Fig 2C).

**Fig 2.**
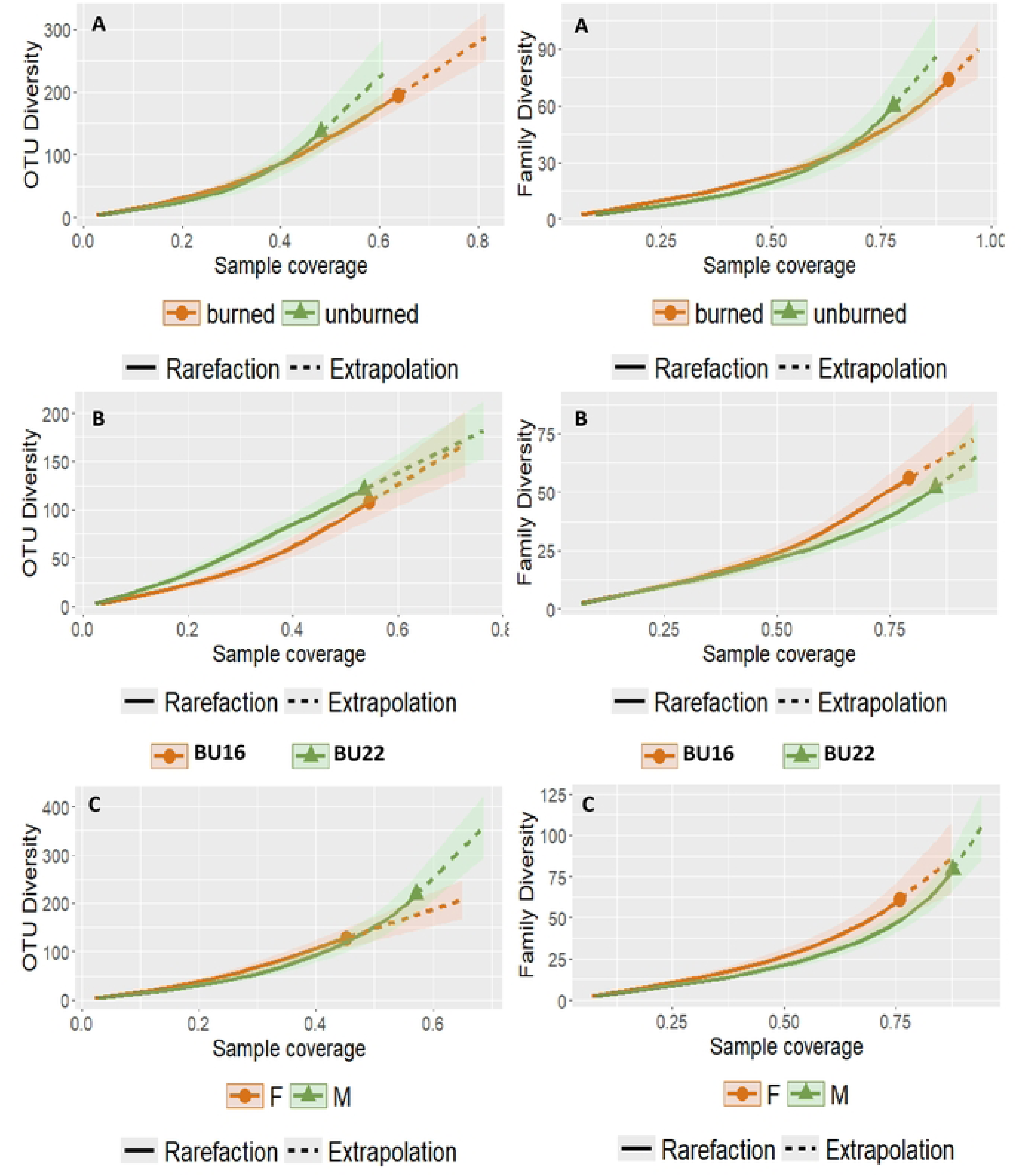
Diet rarefaction curves for (A) fire type (burned, unburned), (B) burned areas (BU16, BU22) and (C) sex (F, female; M, male), showing the observed (full line) and estimated (dashed line) OTU and Family richness, and their respective 84% confidence intervals by sample coverage.

The rank abundance plot (S5 Fig) shows that the most abundant species present in both burned areas is the terrestrial amphipod *Talitroides topitotum*, while in the unburned areas the most abundant OTU is “Arachnida 2”, followed by *T. topitotum*. The BU22 line shows a steep drop after the first most abundant OTU, indicating high dominance by a few species. When compared to BU16, it is possible to infer that BU16 has a more gradual decline in abundance compared to BU22, indicating change in species composition over time.

The PERMANOVA results indicated that Fire type significantly influenced the OTU community composition (F = 1.4672, p = 0.014), while Sex and SVL did not have significant effects (Table 1). The homogeneity of dispersion test confirms that this is not due to unequally dispersed values (F = 0.8968, p = 0.4097). The significant differences obtained between the Fire types were due to differences between recently burned (BU22) and unburned areas (UN) (p = 0.004; Table 1), although the differences between the unburned areas (UN) and the areas burned in 2016 (BU16) were nearly significant (p = 0.051; Table 1). The OTUs *Salticus scenicus* and “*Tapinoma* sp.1” contributed the most to the differences found in PERMANOVA between the UN and BU22. At the Family level, the PERMANOVA did not show significant effects of any of the considered factors in diet composition (Table 1).

**Table 1.**
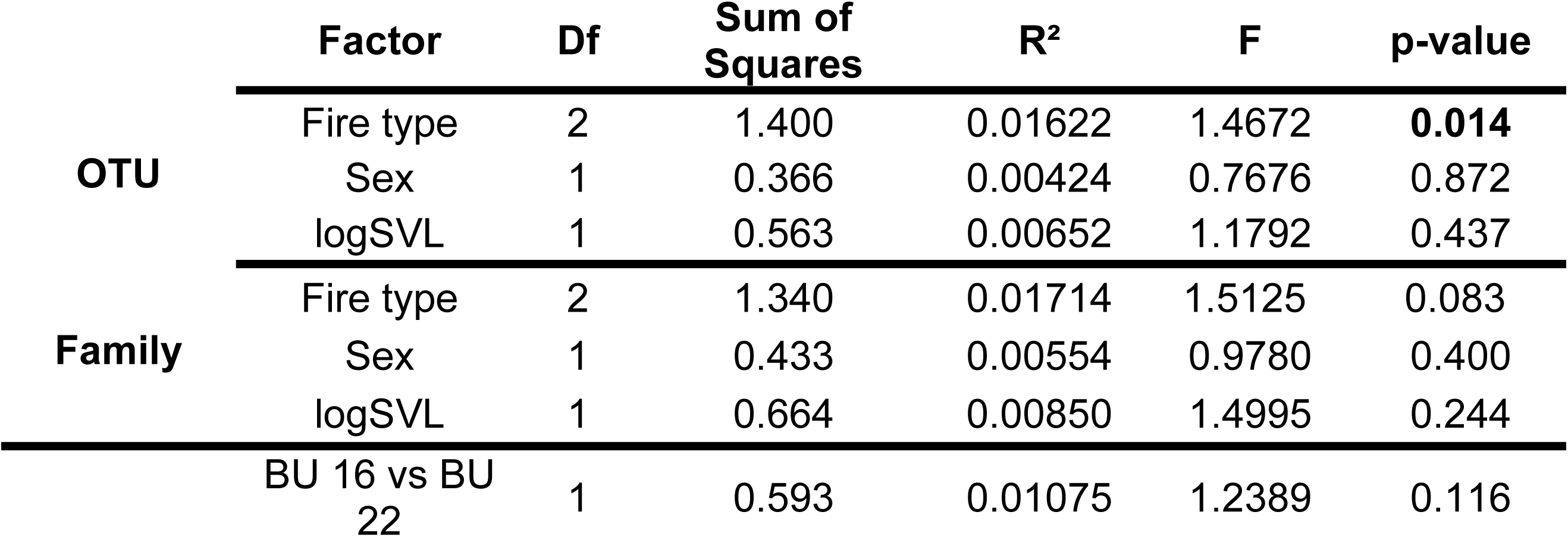
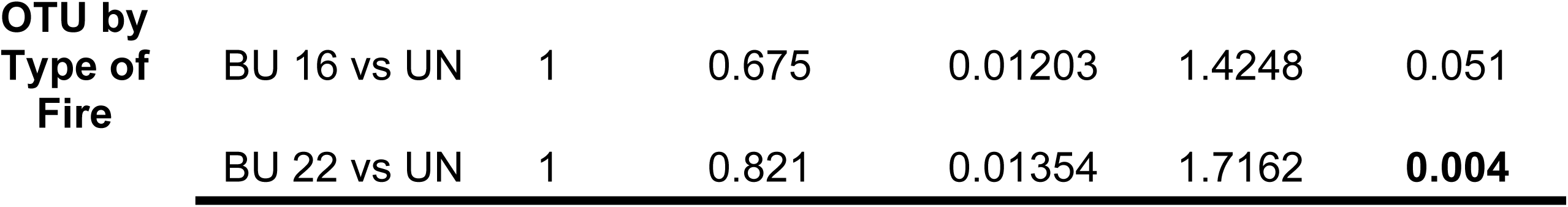
PERMANOVA results for OTU and Family community composition and Pairwise PERMANOVA Comparisons for OTU by Type of Fire. Significant p-values (p < 0.05) are highlighted in bold.

Regarding the RDA, the first two axes accounted for 40.2% and 38.6% of the explained variation, respectively (Fig 3). Fire type showed a statistically significant effect on Family abundance (F = 1.2957, p = 0.045), while sex did not contribute significantly (F = 0.9443, p = 0.546). From the results it was possible to observe specific associations between prey families and fire types: Miridae, Tabanidae and Curculionidae were associated with areas burned in 2016; Cicadellidae and Noctuidae were associated with areas burned in 2022; and Formicidae and Salticidae were associated with unburnt areas. The statistical results for RDA axes indicated that no single axis was individually significant, although the first two axes captured the majority of the constrained variance.

**Fig 3.**
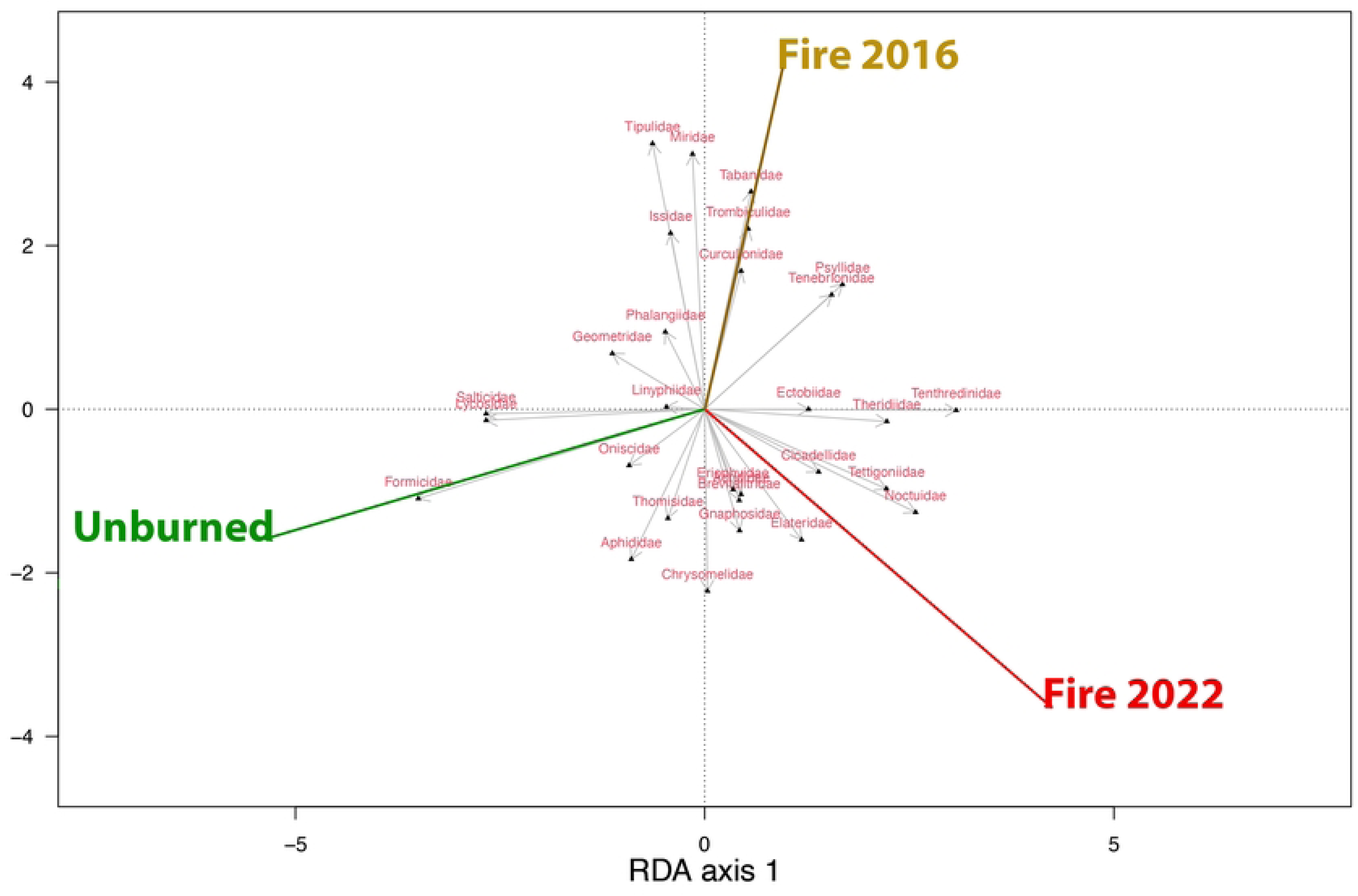
Redundancy analysis (RDA) biplot showing the relationship between prey abundance per Family and fire history. Arrows represent arthropod families, with their orientation and length indicating their contribution to the variation explained by the first two RDA axes (RDA1 and RDA2). Coloured vectors represent plot types: unburned (green), burned in 2016 (yellow) and burned in 2022 (red).

## Discussion

In this study, the effect of wildfires on the trophic niche of *Podarcis lusitanicus* was assessed, by comparing diet richness and composition among specimens captured in sites that were burned in 2022, in 2016, and in unburned areas. Results showed that diet richness does not change significantly between sites and is also not dependent on Sex or SVL. On the other hand, composition was distinct between sites, meaning that fire history influences lizard diets.

Members of the Lacertidae Family are typical examples of generalist species, feeding on a high variety of terrestrial invertebrates [67, 68]. Previous studies on the diet of *P. lusitanicus* have used morphological prey identification through stomach contents [69, 70] or pellets [58]. They revealed that the main prey groups included Hemiptera, Coleoptera, Araneae, Hymenoptera and Diptera. To a lesser extent, Orthoptera, Lepidoptera (including larvae), Opiliones, Miriapoda, Isopoda, Collembola, Diptera larvae, Trichoptera larvae, Mecoptera, Blattodea, Thysanoptera, Dermaptera, Embioptera, as well as vertebrates, Oligochaetes and Gastropods were also identified [69-73]. Our results confirmed that the diet of *P. lusitanicus* is quite diverse, with a taxonomic composition in line with previous morphological studies carried out on this species, highlighting the predominant groups, such as Hemiptera, Coleoptera, Araneae, Hymenoptera and Diptera [74]. However, a far greater diversity of prey items was identified in our study compared to previous ones [70, 74]. Contrary to the genetic approach, diet studies based on morphology tend to underestimate the presence of soft-bodied prey, as they typically only detect items that have not been completely digested [75]. On the other hand, some OTUs identified under DNA metabarcoding may represent secondary predation, which can overestimate prey diversity estimates [76]. Nevertheless, the significant increase in the number of orders identified in this study (26) compared to previous studies using microscopy (15 and 19 in Pérez Mellado (1982) [70] and Carretero et al. (2022) [74], respectively) suggests that the prey spectrum of *P. lusitanicus* is considerably larger than initially reported. In addition, this metabarcoding study provided greater precision in the taxonomic identification of prey, allowing a great number of them to be identified up to the species level (184; see S3 Table), which is usually not possible in studies based on morphological scatology.

### Fire Impact on Richness and Composition

Wildfires in Mediterranean areas cause the fragmentation and destruction of original forest habitats, resulting in a decrease in the size of these areas, increasing the isolation of forest patches and creating a heterogeneous landscape. This creates forest archipelagos dispersed in matrices of open or shrubby vegetation, with different post-wildfire ages [9]. In these open landscapes both vegetation and fauna recover through ecological succession, from low xeric shrubs to tall evergreen forests [77-79]. As a consequence, differences in prey items between areas affected and unaffected by fire would be expected [80]. Essentially, the recovery of prey communities, such as arthropods, after major disturbances like wildfires, can be attributed to two main factors: their resistance to fire, which is their ability to survive when flames pass through, and their resilience, which includes their capacity to recolonize burned areas, take advantage of new resources created by the fire, and adapt as the habitat gradually recovers to its original state [41, 81, 82]. Refugia not only increases immediate survival during a fire but can also facilitate the persistence and recovery of populations in affected areas. In the long term, as vegetation regenerates, these refugia can help re-establish populations, acting as sources of expansion within the burned area or facilitating the arrival of new individuals from outside that area, providing both short-term (food and shelter) and long-term (permanent habitat) resources [81, 83, 84].

Our results demonstrate that lizards in burned areas seem to be able to maintain levels of dietary diversity. This can be explained by the ability of these lizards to adjust their diet, including invertebrate species that are more resilient to fire or those that quickly colonise habitats after the fire. Lacertid lizards are known to show remarkable dietary flexibility in response to environmental changes. Although their diet composition is not determined solely by prey availability [33], they can adapt their diet based on the type of habitat and resources available after a disturbance [68].

In this study, we demonstrate that depending on the fire regime, prey composition is significantly different, corroborating the previously described trophic behaviour in lizards. Specifically, the differences are observed between areas recently burned in 2022 and unburned areas, i.e. those fire regimes supporting the highest structural differences in the habitat. These differences in diet composition were mainly due to the presence of the ant *Tapinoma* sp.1 and the spider *Salticus scenicus* in unburned areas, and their total absence in the 2022 burned sites. The results of the RDA suggest that other prey types also contribute to dietary differences among lizards from different fire types, indicating a more complex pattern of prey availability and selection. The absence of significant differences in diet composition at the Family level suggests that, although fires can alter the composition of specific OTUs, the broader taxonomic structure (e.g., consumption of ants, spiders, etc.) remains relatively stable. This could possibly happen due to the resilience of diversity at the Family level or functional redundancy within these families, meaning that different species within a family can perform similar functions in the ecosystem (as was previously found for bees in Moretti et al., 2009 [85]).

The finding that neither diet richness nor composition differ significantly between males and females suggests that both sexes have similar access to food resources and use them in a similar way, possibly due to a shared ecological niche or similar adaptive behaviours. This is somewhat unexpected, given that prey selection can be influenced by morphological characteristics. Kaliontzopoulou et al (2012) [58] observed that males selected larger and harder prey items, a tendency that correlates positively with head size and bite force, both bigger in males than in females. Considering that the studied populations of *P. lusitanicus* show sexual dimorphism for both size and weight, differences in prey composition were expected between both sexes. However, in dietary assessments using metabarcoding, faecal samples are crushed during DNA extraction, and so size and hardness of the prey are impossible to be determined, which are crucial factors in prey selection by lizards (e.g. Díaz & Carrascal, 1990 [86]). Hence, we are unable to verify whether, despite consuming similar taxonomical groups of arthropods, if males and females of *P. lusitanicus* are selecting specimens of different sizes or even from distinct ontogenic stages.

### Fire Responsive Taxa

Some studies have shown that certain fire-sensitive species undergo a significant decline in their populations soon after a fire (e.g., Santos et al., 2016 [11]). However, other studies indicate that it is rare for wildfires to cause the complete extinction of these species in the affected areas. The chance of an individual immediately surviving a fire depends on the intensity of the fire, the proximity of safe shelters and the behaviours the organism can adopt to avoid the flames and direct heat [83, 87, 88]. The loss of the litter layer and the consequent increase in soil exposure reduce moisture and the amount of organic matter, essential resources for arthropods that depend on them for food and shelter [89]. In addition, increased temperatures in burnt areas can alter the life cycles of arthropods, leading to events such as the early hatching of species, such as grasshoppers [90]. Leaf litter removal also disrupts habitat structure, forcing some species to migrate to unburnt areas in search of better resources [91, 92]. However, the early post-fire succession enhances open areas and promotes a change in dominant animal species, which frequently results in different assemblages in burnt and unburnt areas [93, 94]. Within the burnt plots, species more characteristically from open areas were expected to be found, including those that construct nests, forage in/on the soil, and disperse over longer distances. For unburnt plots, species more typically found in vegetated habitats were expected, especially those with shorter dispersal distances [95]. Therefore, the effect of fire on species richness is highly variable and depends on habitat type, fire intensity, and the taxonomic group considered. For example, different ant taxa can be classified as “fire-tolerant”, “fire-intolerant” or “fire-neutral” depending on their ability to survive and adapt to fire-disturbed habitats. A meta-analysis by Vasconcelos et al. (2017) [96], which synthesized data from 50 studies conducted in various ecosystems, found that the impact of fire on ant diversity varied significantly across vegetation types. The effect was particularly negative in environments where fire is infrequent, while it was minimal or negligible in fire-prone ecosystems. The presence of common ant species, such as *Tetramorium caespitum*, in both burnt and unburnt habitats highlights their high ecological plasticity, although certain species, like the arboreal *Camponotus ruber*, show a more limited range of habitat tolerance [20]. Interestingly, previous studies, such as EL Khayati et al. (2023) [20], reported changes in ant species observed in burnt and unburned areas, and the findings of this study showed that lizards in unburned plots consumed a higher diversity of Formicidae. Similarly, Salticidae (jumping spiders) were also predominantly foraged on by lizards in unburnt habitats. Despite this association, spiders and ants are often considered moderately fire-resistant due to traits that enhance their survival, such as burrowing behaviours, the ability to utilize refuges like logs and rock piles, and their capacity to adapt to post-fire environments [89, 97]. However, the association of these taxa with unburnt areas observed in this study does not necessarily indicate a decrease in their abundance in burnt areas, since this research did not examine pre- and post-fire arthropod diversity directly.

The consumption of several prey types in lizards from burned plots (a result supported by the RDA) suggests that lizards’ diet mirrors availability, and availability is linked to vegetation associated to particular fire type. Thus, families such as Cicadellidae (Hemiptera) and Tettigoniidae (Orthoptera) were frequently more abundant in burned areas [98, 99]. This trend is confirmed in our results, as Cicadellidae and Tettigoniidae are often associated with the regrowth of early successional plants that dominate post-fire environments. The regrowth provides abundant resources such as new leaves for Cicadellidae and open spaces with fresh vegetation for Tettigoniidae. In plots burned in 2016, the families Tabanidae, Miridae, and Curculionidae were strongly associated with post-fire transitional vegetation. Tabanidae, as pollinators, and Miridae and Curculionidae, as plant feeders, likely benefit from the increased availability of nectar and plant material in regenerating landscapes [98, 100]. Notably, Miridae has been reported to maintain high abundances in both burned and unburned sites, likely due to their adaptability to a wide range of vegetation [96, 98].

The dominance of *Talitroides topitotum* in both burned areas suggests that this species may be a fire tolerant species. *Talitroides topitotum* is a terrestrial amphipod native to Asia, widely found in subtropical and temperate regions around the world [101]. The species has been introduced to countries such as Costa Rica [101, 102], Brazil [103] and Indonesia [104] and is able to adapt to diverse temperature and humidity conditions [101]. Due to its high capacity for dispersal and establishment in new environments, *T. topitotum* can become potentially invasive in various habitats, posing a threat to native amphipod species [101, 104]. *Talitroides topitotum* also seems to be associated with areas that were reforested with *Eucalyptus* spp., a culture that covers extensive areas in Brazil [105] and that represents the most widespread tree species in Portuguese mainland [106]. To the best of our knowledge, while this species has not yet been documented in mainland Portugal, it has been recorded in Madeira and Azores archipelagos [107]. These results highlight both the metabarcoding technique and dietary studies of lizards as possible tools in the detection of unrecorded alien arthropod species (e.g. Martins et al, 2022 [108]).

### Diet Studies to assess Arthropod’s Diversity

Ultimately, dietary studies can serve as a tool for assessing arthropod diversity, with reptiles as natural samplers. Traditionally, arthropod diversity studies have used methods such as Malaise traps, flight-intercept traps and pan traps, or Berlese funnels [109]. However, there are studies assessing the diversity of arthropods by analysing the diet of their predators, such as birds [110], and bats [111]. DNA metabarcoding stands out as a powerful tool for identifying prey taxa, offering greater sensitivity and taxonomic resolution compared to morphological analyses [111]. Although this was not the primary aim of our study, by investigating the diet of lizards we obtained data that could be valuable for understanding the diversity of arthropods in Northern Portugal, identifying species not previously recorded and updating the available databases. Validating new localities across a wide selection of invertebrates is complex, since public databases often contain taxonomic errors and geographical uncertainties, which can lead to overestimation or underestimation of species richness in certain areas [112]. Local databases are invaluable in this context, as they can provide more accurate and comprehensive information [113]. For example, while many species were not identified using the Global Biodiversity Information Facility (GBIF, http://www.gbif.org/), specialist sites for Portugal such as Naturdata (https://naturdata.com/) and Lusoborboletas (https://www.lusoborboletaspt.com/), allowed us to greatly refine the number of potentially new records for Portugal. In our study, several prey species such as *Acanthogethes fuscus, Diplazon tetragonus, Formica selysi, Monophadnoides ruficruris* and *Psilopa marginella*, appear to be potentially new records for Portugal, and this information can be used to update the relevant faunistic databases. Swengel (2001) [114] observed that highly mobile insects, including flying species, are typically the first to recolonize burnt landscapes. This mobility is a shared characteristic among most of these species, with the exception of *A. fuscus*.

## Conclusion

This study allowed us to shed new light on the effects of wildfires on the diet of several populations of *Podarcis lusitanicus* lizards in a region regularly affected by these events, such as Northern Portugal. This study suggests that wildfires do not seem to affect the diet richness of *P. lusitanicus* but instead influence the composition of consumed prey items. Specifically, differences are observed between recently burned and unburned habitats, while areas burned 8 years ago and unburned do not show differences in arthropod composition. These results evidence not only the capacity of this species to adapt to this specific disturbance, but also of the habitats’ regenerative power. Ultimately, the results of this study also demonstrate that not only the use of DNA metabarcoding offers a promising window to unravel the interactions between prey and predator with more precision than morphological approaches but also allows to improve the arthropod records. This is particularly important for detecting invasive or crop pest species. In this context, *P. lusitanicus* and other lizard species could act as early indicators of the presence of invasive invertebrate species, underscoring the broader ecological applications of this methodology.

## Acknowledgments

This study was conducted as part of the MSc thesis of Catarina Simões, under the supervision of D. James Harris and Catarina Rato.

We thank Joaquim Faria and Giulia Simbola who participated in sample collection efforts on behalf of this project, Sonia Ferreira for advice about the distribution of invertebrates from Portugal, and Brahim Chergui for the scripts provided.

## Supporting Information

**S1 Table. Number of lizards examined at each site, followed by the fire regime: UN (unburned), BU16 (burned in 2016), and BU22 (burned in 2022).**

**S2 Table. ANOVA results for the effect of Fire type (Locality as a random factor) and Sex on logSVL and logweight. Significant p-values (p < 0.05) are highlighted in bold.**

**S3 Table. GLMM results for Fire type, Sex and SVL (logSVL) on OTU and Family richness in *Podarcis lusitanicus*.**

**S4 Table. Dietary data from *Podarcis lusitanicus*’s.**

**S5 Table. Frequency of occurrence of each OTU prey items present in the diet of *Podarcis lusitanicus* from areas burned in 2016 (BU16), burned in 2022 (BU22) and unburned (UN).**

**S6 Table. Frequency of occurrence of families of prey items present in the diet of *Podarcis lusitanicus* from areas burned in 2016 (BU16), burned in 2022 (BU22) and unburned areas (UN).**

**S1 Fig. Boxplot showing the differences between sexes regarding the logSVL (A) and log Weight (B) across the different types of fire regimes - areas burned in 2016 (BU16), burned in 2022 (BU22) and unburned areas (UN).**

**S2 Fig. Linear regression of log-transformed SVL (logSVL) against log-transformed Weight (logWeight) for female (F) and male (M) individuals.**

**S3 Fig. Histogram representing the frequency of occurrence of the arthropod orders (total of 26) consumed.**

**S4 Fig. Estimated model coefficients and 95% confidence intervals (CIs) of the predictors used to model OTU and Family richness.** Predictor variables had a significant effect when 95% confidence interval did not overlap 0. Areas burned in 2022 (BU22), and unburned areas (UNB) are compared with areas burned in 2016 regarding the type of fire.

**S5 Fig. Rank abundance plot for the 10 more abundant OTUs found in *P. lusitanicus*’ diet, among the different fire types – areas burned in 2016 (BU16), burned in 2022 (BU22) and unburned areas (UN).**

## References

1. Turner MG. Disturbance and landscape dynamics in a changing world. Ecology. 2010;91(10):2833–49.

2. Chuvieco E, Giglio L, Justice C. Global characterization of fire activity: toward defining fire regimes from Earth observation data. Global change biology. 2008;14(7):1488–502.

3. Catry FX, Rego FC, Bação FL, Moreira F. Modeling and mapping wildfire ignition risk in Portugal. International Journal of Wildland Fire. 2009;18(8):921.

4. Hosseini M, Keizer JJ, Pelayo OG, Prats SA, Ritsema C, Geissen V. Effect of fire frequency on runoff, soil erosion, and loss of organic matter at the micro-plot scale in north-central Portugal. Geoderma. 2016;269:126–37.

5. Keeley JE, Bond WJ, Bradstock RA, Pausas JG, Rundel PW. Fire in Mediterranean ecosystems: ecology, evolution and management: Cambridge University Press; 2011.

6. Pausas J, Verdú M. Plant persistence traits in fire-prone ecosystems of the Mediterranean basin: a phylogenetic approach. Oikos. 2005;109(1):196–202.

7. Syphard AD, Radeloff VC, Hawbaker TJ, Stewart SI. Conservation threats due to human-caused increases in fire frequency in Mediterranean-climate ecosystems. Conservation Biology. 2009;23(3):758–69.

8. Izhaki I. The impact of fire on vertebrates in the Mediterranean Basin: an overview. Israel Journal of Ecology and Evolution. 2012;58(2-3):221–33.

9. Sarà M, Bellia E, Milazzo A. Fire disturbance disrupts co-occurrence patterns of terrestrial vertebrates in Mediterranean woodlands. Journal of Biogeography. 2006;33(5):843–52.

10. Jolly CJ, Dickman CR, Doherty TS, van Eeden LM, Geary WL, Legge SM, et al. Animal mortality during fire. Global Change Biology. 2022;28(6):2053–65.

11. Santos X, Badiane A, Matos C. Contrasts in short-and long-term responses of Mediterranean reptile species to fire and habitat structure. Oecologia. 2016;180:205–16.

12. Santos X, Chergui B, Belliure J, Moreira F, Pausas JG. Reptile responses to fire across the western Mediterranean Basin. Conservation Biology. 2024:e14326.

13. Moyo S. Community Responses to Fire: A Global Meta-Analysis Unravels the Contrasting Responses of Fauna to Fire. Earth. 2022;3(4):1087–111.

14. Sutherland EF, Dickman CR. Mechanisms of recovery after fire by rodents in the Australian environment: a review. Wildlife Research. 1999;26(4):405–19.

15. Nieman WA, van Wilgen BW, Radloff FG, Leslie AJ. A review of the responses of medium-to large-sized African mammals to fire. African Journal of Range & Forage Science. 2022;39(3):249–63.

16. Ferrenberg S, Wickey P, Coop JD. Ground-dwelling arthropod community responses to recent and repeated wildfires in conifer forests of northern New Mexico, USA. Forests. 2019;10(8):667.

17. Certini G, Moya D, Lucas-Borja ME, Mastrolonardo G. The impact of fire on soil-dwelling biota: A review. Forest Ecology and Management. 2021;488:118989.

18. Karpestam E, Merilaita S, Forsman A. Reduced predation risk for melanistic pygmy grasshoppers in post-fire environments. Ecology and Evolution. 2012;2(9):2204–12.

19. Mola JM, Miller MR, O’Rourke SM, Williams NM. Wildfire reveals transient changes to individual traits and population responses of a native bumble bee *Bombus vosnesenskii*. Journal of Animal Ecology. 2020;89(8):1799–810.

20. EL Khayati M, Chergui B, Barranco P, Fahd S, Ruiz JL, Taheri A, et al. Assessing the Response of Different Soil Arthropod Communities to Fire: A Case Study from Northwestern Africa. Fire. 2023;6(5):206.

21. Santos X, Mateos E, Bros V, Brotons L, De Mas E, Herraiz JA, et al. Is response to fire influenced by dietary specialization and mobility? A comparative study with multiple animal assemblages. PLoS One. 2014;9(2):e88224.

22. Doherty TS, Geary WL, Jolly CJ, Macdonald KJ, Miritis V, Watchorn DJ, et al. Fire as a driver and mediator of predator–prey interactions. Biological Reviews. 2022;97(4):1539–58.

23. Hoare S. The possible role of predator–prey dynamics as an influence on early hominin use of burned landscapes. Evolutionary Anthropology: Issues, News, and Reviews. 2019;28(6):295–302.

24. Alemany I, Pérez-Cembranos A, Pérez-Mellado V, Castro JA, Picornell A, Ramon C, et al. DNA metabarcoding the diet of *Podarcis* lizards endemic to the Balearic Islands. Current Zoology. 2022:zoac073.

25. Mateus P, Fernandes PM. Forest fires in Portugal: dynamics, causes and policies. Forest context and policies in Portugal: present and future challenges. 2014:97–115.

26. San-Miguel-Ayanz J, Oom D, Artes T, Viegas DX, Fernandes P, Faivre N, et al. Forest fires in Portugal in 2017. Science for disaster risk management. 2020:413–30.

27. Ferreira D, Pinho C, Brito JC, Santos X. Increase of genetic diversity indicates ecological opportunities in recurrent-fire landscapes for wall lizards. Scientific Reports. 2019;9(1):5383.

28. Meira Castro AC, Nunes A, Sousa A, Lourenço L. Mapping the causes of forest fires in portugal by clustering analysis. Geosciences. 2020;10(2):53.

29. Pausas JG, Llovet J, Rodrigo A, Vallejo R. Are wildfires a disaster in the Mediterranean basin?–A review. International Journal of wildland fire. 2008;17(6):713–23.

30. Nunes AN. Regional variability and driving forces behind forest fires in Portugal an overview of the last three decades (1980–2009). Applied Geography. 2012;34:576–86.

31. Carretero MA. An integrated assessment of a group with complex systematics: the Iberomaghrebian lizard genus *Podarcis* (Squamata, Lacertidae). Integrative Zoology. 2008;3(4):247–66.

32. Harris DJ, Sa-Sousa P. Molecular phylogenetics of Iberian wall lizards (*Podarcis*): is *Podarcis hispanica* a species complex? Molecular Phylogenetics and Evolution. 2002;23(1):75–81.

33. Carretero MA. From set menu to a la carte. Linking issues in trophic ecology of Mediterranean lacertids. Italian Journal of Zoology. 2004;71(S2):121-33.

34. Speybroeck J, Beukema W, Bok B, Van Der Voort J. Field guide to the amphibians and reptiles of Britain and Europe: Bloomsbury publishing; 2016.

35. Rato C, Sreelatha LB, Gómez-Ramírez F, Carretero MA. A Pleistocene Biogeography in Miniature: The Small-Scale Evolutionary History of *Podarcis lusitanicus* (Squamata, Lacertidae). Journal of Biogeography. 2025;52(1):186–98.

36. Geniez P, Sa-Sousa P, Guillaume CP, Cluchier A, Crochet P-A. Systematics of the *Podarcis hispanicus* complex (Sauria, Lacertidae) III: valid nomina of the western and central Iberian forms. Zootaxa. 2014;3794(1):1–51.

37. Ferreira D, Žagar A, Santos X. Uncovering the rules of (reptile) species coexistence in transition zones between bioregions. Salamandra. 2017;53(2).

38. Diego-Rasilla FJ, Perez-Mellado V. Home range and habitat selection by *Podarcis hispanica* (Squamata, Lacertidae) in Western Spain. Folia Zoologica-Praha-. 2003;52(1):87–98.

39. Ferreira D, Mateus C, Santos X. Responses of reptiles to fire in transition zones are mediated by bioregion affinity of species. Biodiversity and Conservation. 2016;25:1543–57.

40. Calheiros T, Benali A, Pereira M, Silva J, Nunes J. Drivers of extreme burnt area in Portugal: fire weather and vegetation. Natural Hazards and Earth System Sciences. 2022;22(12):4019–37.

41. Khayati ME, Chergui B, Taheri A, Fahd S, Santos X. Differential response to fire in ground vs. vegetation arthropod communities. Journal of Insect Conservation. 2023;27(4):601–13.

42. Barata M, Harris DJ. Cryptic variation in the Moroccan high altitude lizard *Atlantolacerta andreanskyi* (Squamata: Lacertidae). African Journal of Herpetology. 2015;64(1):1–17.

43. Perera A, Pérez-Mellado V, Carretero MA, Harris DJ. Variation between populations in the diet of the Mediterranean lizard *Lacerta perspicillata*. The Herpetological Journal. 2006;16(2):107–13.

44. Sá-Sousa P, Vicente L, Crespo E. Morphological variability of *Podarcis hispanica* (Sauria: lacertidae) in Portugal. Amphibia-Reptilia. 2002;23(1):55–69.

45. Maudet C, Luikart G, Dubray D, Von Hardenberg A, Taberlet P. Low genotyping error rates in wild ungulate faeces sampled in winter. Molecular Ecology Notes. 2004;4(4):772–5.

46. Vamos EE, Elbrecht V, Leese F. Short COI markers for freshwater macroinvertebrate metabarcoding. PeerJ Preprints; 2017. Report No.: 2167-9843.

47. Elbrecht V, Braukmann TW, Ivanova NV, Prosser SW, Hajibabaei M, Wright M, et al. Validation of COI metabarcoding primers for terrestrial arthropods. PeerJ. 2019;7:e7745.

48. Tournayre O, Leuchtmann M, Filippi-Codaccioni O, Trillat M, Piry S, Pontier D, et al. In silico and empirical evaluation of twelve metabarcoding primer sets for insectivorous diet analyses. Ecology and Evolution. 2020;10(13):6310–32.

49. Zhang J, Kobert K, Flouri T, Stamatakis A. PEAR: a fast and accurate Illumina Paired-End reAd mergeR. Bioinformatics. 2014;30(5):614–20.

50. Boyer F, Mercier C, Bonin A, Le Bras Y, Taberlet P, Coissac E. obitools: A unix-inspired software package for DNA metabarcoding. Molecular Ecology Resources. 2016;16(1):176–82.

51. Rognes T, Flouri T, Nichols B, Quince C, Mahé F. VSEARCH: a versatile open source tool for metagenomics. PeerJ. 2016;4:e2584.

52. Mata VA, Amorim F, Corley MF, McCracken GF, Rebelo H, Beja P. Female dietary bias towards large migratory moths in the European free-tailed bat (*Tadarida teniotis*). Biology Letters. 2016;12(3):20150988.

53. Evans HK, Bunch AJ, Schmitt JD, Hoogakker FJ, Carlson KB. High-throughput sequencing outperforms traditional morphological methods in Blue Catfish diet analysis and reveals novel insights into diet ecology. Ecology and Evolution. 2021;11(10):5584–97.

54. Frøslev TG, Kjøller R, Bruun HH, Ejrnæs R, Brunbjerg AK, Pietroni C, et al. Algorithm for post-clustering curation of DNA amplicon data yields reliable biodiversity estimates. Nature communications. 2017;8(1):1188.

55. Buchner D, Leese F. BOLDigger–a Python package to identify and organise sequences with the Barcode of Life Data systems. Metabarcoding and Metagenomics. 2020;4:e53535.

56. Kearse M, Moir R, Wilson A, Stones-Havas S, Cheung M, Sturrock S, et al. Geneious Basic: an integrated and extendable desktop software platform for the organization and analysis of sequence data. Bioinformatics. 2012;28(12):1647–9.

57. Costa GC, Vitt LJ, Pianka ER, Mesquita DO, Colli GR. Optimal foraging constrains macroecological patterns: body size and dietary niche breadth in lizards. Global Ecology and Biogeography. 2008;17(5):670–7.

58. Kaliontzopoulou A, Adams DC, van der Meijden A, Perera A, Carretero MA. Relationships between head morphology, bite performance and ecology in two species of *Podarcis* wall lizards. Evolutionary Ecology. 2012a;26:825–45.

59. Russell L. Emmeans: estimated marginal means, aka least-squares means. R package version. 2018;1(2).

60. R Core Team R. R: A language and environment for statistical computing. R foundation for statistical computing Vienna, Austria; 2013.

61. Hsieh T, Ma K, Chao A. iNEXT: an R package for rarefaction and extrapolation of species diversity (H ill numbers). Methods in ecology and evolution. 2016;7(12):1451–6.

62. MacGregor-Fors I, Payton ME. Contrasting diversity values: statistical inferences based on overlapping confidence intervals. PloS one. 2013;8(2):e56794.

63. Chao A, Jost L. Coverage-based rarefaction and extrapolation: standardizing samples by completeness rather than size. Ecology. 2012;93(12):2533–47.

64. Wickham H. ggplot2: Elegant Graphics for Data Analysis. New York, NY, USA: Springer-Verlag; 2016.

65. Oksanen J, Simpson G, Blanchet F, Kindt R, Legendre P, Minchin P, et al. vegan: Community Ecology Package. R package version 2.6–2. 2022. Google Scholar There is no corresponding record for this reference. 2022.

66. Martinez Arbizu P. pairwiseAdonis: Pairwise multilevel comparison using adonis. R package version 04. 2020;1.

67. Arnold EN. Resource partition among lacertid lizards in southern Europe. Journal of Zoology. 1987;1(4):739–82.

68. Sagonas K, Pafilis P, Lymberakis P, Donihue CM, Herrel A, Valakos ED. Insularity affects head morphology, bite force and diet in a Mediterranean lizard. Biological Journal of the Linnean Society. 2014;112(3):469–84.

69. Bas S. La comunidad herpetológica de Caurel: biogeografía y ecología. Amphibia-Reptilia. 1982;3(1):1–26.

70. Pérez Mellado V. Estructura en una taxocenosis de Lacertidae (Sauria, Reptilia) del sistema central. Mediterránea Serie de Estudios Biológicos, N 6 (diciembre 1982); pp 39-64. 1982.

71. Braña F. Biogeografía, biología y estructura de nichos de la taxocenosis de saurios de Asturias. Unpubl PhD Thesis Univ Oviedo Oviedo, Spain. 1984.

72. Galán P. Anfibios y reptiles del Parque Nacional de las Islas Atlánticas de Galicia: faunística, biología y conservación: Ministerio de Medo Ambiente, Secretaría General de Medio Ambiente, Organismo Autónomo Parques Nacionales.; 2003.

73. Kaliontzopoulou A, Carretero MA, Llorente GA. Morphology of the *Podarcis* wall lizards (Squamata: Lacertidae) from the Iberian Peninsula and North Africa: patterns of variation in a putative cryptic species complex. Zoological Journal of the Linnean Society. 2012b;164(1):173–93.

74. Carretero MÁ, Galán P, Salvador Milla A. Lagartija lusitana–Podarcis lusitanicus Geniez, Sá-Sousa, Guillaume, Cluchier y Crochet, 2014. 2022.

75. Ingerson-Mahar J. Relating diet and morphology in adult carabid beetles. In: J H, editor. The agroecology of carabid beetles. Andover, UK: Intercept; 2002. p. 111-36.

76. da Silva LP, Mata VA, Lopes PB, Pereira P, Jarman SN, Lopes RJ, et al. Advancing the integration of multi-marker metabarcoding data in dietary analysis of trophic generalists. Molecular Ecology Resources. 2019;19(6):1420–32.

77. Katsimanis N, Dretakis M, Akriotis T, Mylonas M. Breeding bird assemblages of eastern Mediterranean shrublands: composition, organisation and patterns of diversity. Journal of Ornithology. 2006;147:419–27.

78. Moreira F, Delgado A, Ferreira S, Borralho R, Oliveira N, Inácio M, et al. Effects of prescribed fire on vegetation structure and breeding birds in young *Pinus pinaster* stands of northern Portugal. Forest Ecology and Management. 2003;184(1-3):225–37.

79. Suárez-Seoane S, Osborne PE, Baudry J. Responses of birds of different biogeographic origins and habitat requirements to agricultural land abandonment in northern Spain. Biological Conservation. 2002;105(3):333–44.

80. Viljur ML, Abella SR, Adámek M, Alencar JBR, Barber NA, Beudert B, et al. The effect of natural disturbances on forest biodiversity: an ecological synthesis. Biological Reviews. 2022;97(5):1930–47.

81. Robinson NM, Leonard SW, Ritchie EG, Bassett M, Chia EK, Buckingham S, et al. Refuges for fauna in fire-prone landscapes: their ecological function and importance. Journal of Applied Ecology. 2013;50(6):1321–9.

82. Steinitz O, Shohami D, Ben-Shlomo R, Nathan R. Genetic consequences of fire to natural populations. Israel Journal of Ecology and Evolution. 2012;58(2-3):205–20.

83. Banks SC, Dujardin M, McBurney L, Blair D, Barker M, Lindenmayer DB. Starting points for small mammal population recovery after wildfire: recolonisation or residual populations? Oikos. 2011;120(1):26–37.

84. Watson S, Taylor R, Nimmo D, Kelly L, Clarke M, Bennett A. The influence of unburnt patches and distance from refuges on post-fire bird communities. Animal Conservation. 2012;15(5):499–507.

85. Moretti M, De Bello F, Roberts SP, Potts SG. Taxonomical vs. functional responses of bee communities to fire in two contrasting climatic regions. Journal of Animal Ecology. 2009:98–108.

86. Díaz JA, Carrascal LM. Prey size and food selection of *Psammodromus algirus* (Lacertidae) in central Spain. Journal of Herpetology. 1990:342–7.

87. Kelly LT, Brotons L, Giljohann KM, McCarthy MA, Pausas JG, Smith AL. Bridging the divide: integrating animal and plant paradigms to secure the future of biodiversity in fire-prone ecosystems. Fire. 2018;1(2):29.

88. Whelan R, Rodgerson L, Dickman C, Sutherland E. Critical life cycles of plants and animals: developing a process-based understanding of population changes in fire-prone landscapes. In: Bradstock RA WJ, Gill AM, editor. Flammable Australia: The fire regimes and biodiversity of a continen. Cambridge: Cambridge University Press; 2002. p. 94-124.

89. Podgaiski LR, Joner F, Lavorel S, Moretti M, Ibanez S, Mendonça Jr MdS, et al. Spider trait assembly patterns and resilience under fire-induced vegetation change in South Brazilian grasslands. PloS one. 2013;8(3):e60207.

90. Meyer C, Whiles M, Charlton R. Life history, secondary production, and ecosystem significance of acridid grasshoppers in annually burned and unburned tallgrass prairie. American Entomologist. 2002;48(1):52–61.

91. Parmenter RR, Kreutzian M, Moore DI, Lightfoot DC. Short-term effects of a summer wildfire on a desert grassland arthropod community in New Mexico. Environmental entomology. 2011;40(5):1051–66.

92. Riechert SE, Reeder WG, editors. Effects of fire on spider distribution in southwestern Wisconsin prairies. Proceedings of the Second Midwest Prairie Conference; 1972: Madison, Wisconsin. p.75-90.

93. Apigian KO, Dahlsten DL, Stephens SL. Fire and fire surrogate treatment effects on leaf litter arthropods in a western Sierra Nevada mixed-conifer forest. Forest Ecology and Management. 2006;221(1-3):110–22.

94. Moretti M, Obrist MK, Duelli P. Arthropod biodiversity after forest fires: winners and losers in the winter fire regime of the southern Alps. Ecography. 2004;27(2):173–86.

95. Vidal-Cordero JM, Arnan X, Rodrigo A, Cerdá X, Boulay R. Four-year study of arthropod taxonomic and functional responses to a forest wildfire: Epigeic ants and spiders are affected differently. Forest Ecology and Management. 2022;520:120379.

96. Vasconcelos HL, Maravalhas JB, Cornelissen T. Effects of fire disturbance on ant abundance and diversity: a global meta-analysis. Biodiversity and Conservation. 2017;26(1):177–88.

97. Pressler Y, Moore JC, Cotrufo MF. Below ground community responses to fire: meta-analysis reveals contrasting responses of soil microorganisms and mesofauna. Oikos. 2019;128(3):309–27.

98. Cancelado R, Yonke TR. Effect of prairie burning on insect populations. Journal of the Kansas Entomological Society. 1970:274–81.

99. Nagel HG. Effect of spring prairie burning on herbivorous and non-herbivorous arthropod populations. Journal of the Kansas Entomological Society. 1973:485–96.

100. Andersen AN. Responses of ground-foraging ant communities to three experimental fire regimes in a savanna forest of tropical Australia. Biotropica. 1991:575–85.

101. Umaña-Castro R, Cambronero-Granados JA, Carvajal-Sánchez JP, Alfaro-Montoya J. Identificación molecular y distribución potencial del anfípodo terrestre *Talitroides topitotum* (Crustacea: Amphipoda: Talitridae) en Costa Rica. Acta Biológica Colombiana. 2018;23(1):104–15.

102. Alfaro-Montoya J, Umaña Castro R. Primer registro e histología básica del anfípodo terrestre *Talitroides topitotum* (Amphipoda: Talitridae), introducido en las zonas montañosas de Heredia, Costa Rica. Archivos. 5: Universidad Estatal a Distancia (Costa Rica); 2013.

103. Brisotto G, Ayres-Peres L, Santos S. Estrutura populacional de *Talitroides topitotum* (Crustacea: Amphipoda: Talitridae) em Santa Maria, região central do Rio Grande do Sul, Brasil. Iheringia Série Zoologia. 2022;112:e2022017.

104. Daneliya M, Wowor D. Cosmopolitan landhopper *Talitroides topitotum* (Crustacea, Amphipoda, Talitridae) in Java, Indonesia. Check List. 2016;12(4):1–9.

105. Lopes OL, Masunari S. Biologia reprodutiva de *Talitroides topitotum* (Burt)(Crustacea, Amphipoda, Talitridae) na Serra do Mar, Guaratuba, Paraná, Brasil. Revista brasileira de Zoologia. 2004;21:755–9.

106. Fernandes P, Antunes C, Pinho P, Máguas C, Correia O. Natural regeneration of *Pinus pinaster* and *Eucalyptus globulus* from plantation into adjacent natural habitats. Forest Ecology and Management. 2016;378:91–102.

107. Secretariat G. GBIF Backbone Taxonomy Checklist dataset; 2022. Flammulina ononidis. 2024.

108. Martins B, Silva-Rocha I, Mata VA, Gonçalves Y, Rocha R, Rato C. Trophic interactions of an invasive gecko in an endemic-rich oceanic island: insights using DNA metabarcoding. Frontiers in Ecology and Evolution. 2022;10:1044230.

109. Marshall SA, Anderson R, Roughley R, Behan-Pelletier V, Danks H. Terrestrial arthropod biodiversity: planning a study and recommended sampling techniques. Bulletin of the Entomological Society of Canada. 1994;26(1):33.

110. Muñoz CE, Ippi SG, Celis Diez JL, Salinas D, Armesto JJ. Arthropods in the diet of the bird assemblage from a forested rural landscape in northern Chiloé island, Chile: a quantitative study. Ornitología Neotropical. 2017;28:191-9.

111. Zeale MR, Butlin RK, Barker GL, Lees DC, Jones G. Taxon-specific PCR for DNA barcoding arthropod prey in bat faeces. Molecular Ecology Resources. 2011;11(2):236–44.

112. Maldonado C, Molina CI, Zizka A, Persson C, Taylor CM, Albán J, et al. Estimating species diversity and distribution in the era of B ig D ata: to what extent can we trust public databases? Global Ecology and Biogeography. 2015;24(8):973–84.

113. Meier R, Dikow T. Significance of specimen databases from taxonomic revisions for estimating and mapping the global species diversity of invertebrates and repatriating reliable specimen data. Conservation Biology. 2004;18(2):478–88.

114. Swengel AB. A literature review of insect responses to fire, compared to other conservation managements of open habitat. Biodiversity & Conservation. 2001;10:1141–69.

